# Non-coding RNAs in fluid shear stress-driven and reactive oxygen species-mediated colon cancer metastasis

**DOI:** 10.1101/2020.08.31.275065

**Authors:** Siluveru KrishnaPriya, Satarupa Banerjee, Devarajan Karunagaran, G. K. Suraishkumar

**Affiliations:** Department of Biotechnology, Bhupat and Jyoti Mehta School of Biosciences, IIT Madras, Chennai-600036, India; School of Bioscience and Technology, Vellore Institute of Technology, Vellore

**Keywords:** Colon cancer metastasis, fluid shear stress, reactive oxygen species, microRNAs, long non-coding RNAs, circular RNAs, network analysis

## Abstract

**Background:** Colon adenocarcinoma (COAD) is the third most common cancer in the world. Fluid shear stress (FSS) and intracellular reactive oxygen species (ROS) levels are known to mediate COAD metastasis. The present work was performed to explore the role of regulatory non-coding RNA molecules associated with FSS and ROS in COAD metastasis.

**Methods:** The interactions between the mRNAs associated with FSS and ROS, the corresponding miRNAs, lncRNAs and circRNAs in COAD metastasis were used to generate the mRNA-miRNA-lncRNA-circRNA network. The expression levels of the RNAs in the network were also considered besides the identification of RNA hubs and modules. Further, functional enrichment and survival analysis of the significant miRNAs together with the OncoPrint as well as survival analysis of the selected mRNAs were performed. Subsequently, their functional role was also corroborated with existing literature.

**Results:** Ten significant miRNA hubs were identified, out of which hsa-miR-17-5p and hsa-miR-20a-5p were found to interact with a lncRNA, CCAT2 and hsa-miR-335 was found to interact with four circRNAs. 60% of the FSS and ROS associated mRNAs and 90% of the top 10 miRNA-enriched pathways that emerged from the functional analysis were reported to be involved in COAD metastasis. 15 significant miRNAs were identified in ten different modules suggesting their importance in FSS and ROS mediated COAD metastasis. Finally, ten miRNAs and three mRNAs associated with FSS and/or ROS were identified as significant overall survival markers; 33 mRNAs were also identified as metastasis-free survival markers whereas 15 mRNAs showed >10% gene alterations in TCGA-COAD data and hence emerged as significant molecular markers in the process.

**Conclusion:** We hypothesize that the biologically significant RNAs identified in this integrated analysis may provide valuable insights to understand the molecular mechanism of the FSS driven and ROS mediated COAD metastasis and to design efficient treatment strategies.

## Introduction

Colon adenocarcinoma (COAD) is the third leading cause of cancer-related deaths worldwide and affects nearly 9% and 8% of females and males, respectively, and the 5-year overall survival rate for the patients is only 65% [1]. Though in most of the cases, primary cancer gets cured by chemotherapy, radiation therapy, and/or surgery, more than 90% of cancer-related deaths are caused due to metastasis [2], a process by which the cancer cells migrate from the primary target organ to the secondary organ via the blood flow [2]. Metastasis is enhanced by fluid shear stress (FSS) in the circulating microenvironment directly, or indirectly via the modulation of the intracellular levels of reactive oxygen species (ROS) [3, 4]. Inter-regulatory cross-talks between intracellular ROS levels and the microRNA (miRNA) expression also control metastasis [5]. FSS, ROS, and non-coding RNAs have been studied in cancer cells, but whether FSS has a regulatory role on the expression of non-coding RNAs in metastasis is not known.

FSS is a major physical factor that contributes significantly to drive the cancer metastatic cascade [4, 6]. As soon as the cancer cells enter into the blood circulation, they are exposed to the mechanical force of shear stress, which drives them to undergo apoptosis [4, 7, 8]. However, FSS also upregulates mechanosensitive molecules of the more-resistant circulating tumor cells, alters the inherent molecular properties and enhances their metastatic ability [7]. The biomechanical forces of hemodynamic shear stress also induce ROS/NO generation, promote epithelial-mesenchymal transition (EMT) [9], and in turn, enhance migration and extravasation of surviving tumor cells [3]. ROS acts as a second messenger in several signaling pathways, activates or inhibits various kinases, phosphatases, and transcription factors [5, 10, 11]. Since cancer cells are characterized by higher intracellular ROS levels than the neighboring normal cells, the study of the ROS-mediated metastatic progression may pave a way to find targeted cancer therapy [10]. Though the antioxidant treatment prevents shear stress-triggered, and ROS mediated metastasis [3] in some cases, FSS and treatment with ROS-generating drugs show a synergistic effect and enhance apoptosis of metastatic cells significantly [12]. Therefore, the molecular changes in the circulatory tumor cells mediated by ROS under FSS conditions are crucial to understand metastasis.

In addition to the coding mRNAs, non-coding RNA molecules such as miRNAs and long non-coding RNAs (lncRNAs) are known to play crucial regulatory roles in COAD metastasis [13, 14], but the role of circular RNAs (circRNAs) in the process is not yet much explored.

The miRNAs are ~22 nucleotide non-coding RNAs, capable of degrading or blocking the translation of the protein-coding mRNA targets by binding, mostly, to their 3’UTRs [15]. They regulate multiple mRNAs simultaneously based on sequence complementarity and thus play diverse roles in cellular metabolic processes [15]. Recent studies revealed that lncRNAs, which are >200 nucleotide long molecules derived from intergenic regions, also contribute significantly to COAD metastasis [16, 17] via chromatin remodeling, transcriptional, or post-transcriptional regulation of the target molecules [14, 18]. Among all the human gene transcripts across different cell types, circRNAs are reported as the most important and dominant regulatory molecules [19], whose expression profiles are tightly regulated and specific for the cell/tissue type or developmental stage [20–22]. These comprise a vast group of covalently closed loops of non-coding RNA molecules [23, 24] and are known to exhibit sequence-specific complementary binding to multiple miRNAs and/or RNA binding proteins (RBPs), often at the same time, termed as sponging [24, 25].

Since shear stress affects the regulatory miRNAs in vascular endothelial cells [26–30], we speculate that FSS may cause alterations in expression profiles of miRNAs and other non-coding RNAs in the circulatory tumor cells also, which can probably promote their metastatic ability. Though earlier reports showed some crosstalks between FSS and ROS [3, 12], as well as ROS and miRNA [5], the association between FSS and non-coding RNAs in cancer metastasis is unexplored and thus addressed here. We hypothesize that the regulatory non-coding RNAs play a significant role in the process. We aim to identify the significant RNAs associated with FSS driven and ROS-mediated COAD metastasis, which is expected to contribute toward better insights for more efficient COAD management and therapy.

To investigate the interconnections between FSS-related, ROS-related and non-coding RNAs in COAD metastasis, a novel approach of interactive network analysis was used. A biological network is the visualization of all the interactions between the molecules associated with the various phenomena involved in a physiological process; it provides a platform to track down the pathway and to understand the molecular mechanism by which their biological functions mediate the process [31]. Although different types of molecules in a cell are associated with diverse phenomena possess their specific physiological functions, the combined set of all these interactions (network) and the extent of modulation that results from each of those interactions ultimately determines the fate of that cell [32]. Such complex networks were generated in this study to depict the interactions among different types of RNA molecules associated with FSS and ROS, to elucidate the crosstalk between them and to understand the circulatory microenvironment that mediates COAD metastasis.

## Materials and methods

### Selection of mRNAs from GO and HCMDB, retrieval of their interacting miRNA, lncRNA and circRNA partners, followed by functional enrichment analysis and assessment of their biological significance

Gene lists associated with the “cellular response to fluid shear stress (GO:0034405)” and “reactive oxygen species metabolic process (GO:0072593)” in *Homo sapiens*, were downloaded from the EMBL-EBI Quick Gene Ontology (GO) database [33]. Genes associated with COAD metastasis (experimentally validated), were downloaded from the Human Cancer Metastasis Database (HCMDB) [34]. The differentially expressed (DE) miRNAs in COAD, along with their expression status, were obtained from dbDEMC 2.0 [35]. DE protein-coding genes in COAD with |log2FC (fold change) | 1.0, were identified using LIMMA, from GEPIA [36]. The common genes among FSS, ROS, HCMDB-COAD, and GEPIA-DE-COAD were identified using the Venn diagram plotted in InteractiVenn (Figure 2a) [37]. Interactions among the selected mRNAs were obtained from the starBase v2.0 database [38], and the interactions of the COAD DE miRNAs with mRNAs of FSS, ROS, and HCMDB-COAD separately were obtained using DIANA-TarBase v7.0 [39]. Common COAD DE miRNAs, interacting with the mRNAs associated with FSS, ROS, and HCMDB-COAD together, were also identified using a Venn diagram (Figure 2b). The functional enrichment of these miRNAs identified in the study was performed using miRNA Enrichment Analysis and Annotation webtool (miEAA) [40]. Then, the top 10 most enriched pathways irrespective of pathway types, viz. Reactome or Panther (Figure 2c), the top 5 most significant Reactome pathways, and GO associated with them (with a P-value < 0.05) (Supplementary figure 1a, 1b) were identified. Subsequently, a literature survey was performed using Google Scholar and PubMed to assess the biological significance of all these enriched pathways as well as the FSS/ROS associated genes and their percentage of significance was evaluated. The identification of lncRNA that interacts with the selected COAD-DE-miRNAs was done using DIANA-LncBase v2 [41] and the circRNAs interacting with the selected miRNAs and mRNAs (RBPs) were obtained from CircInteractome [42]. The workflow proposed in the study is provided in Figure 1.

**Figure 1.**
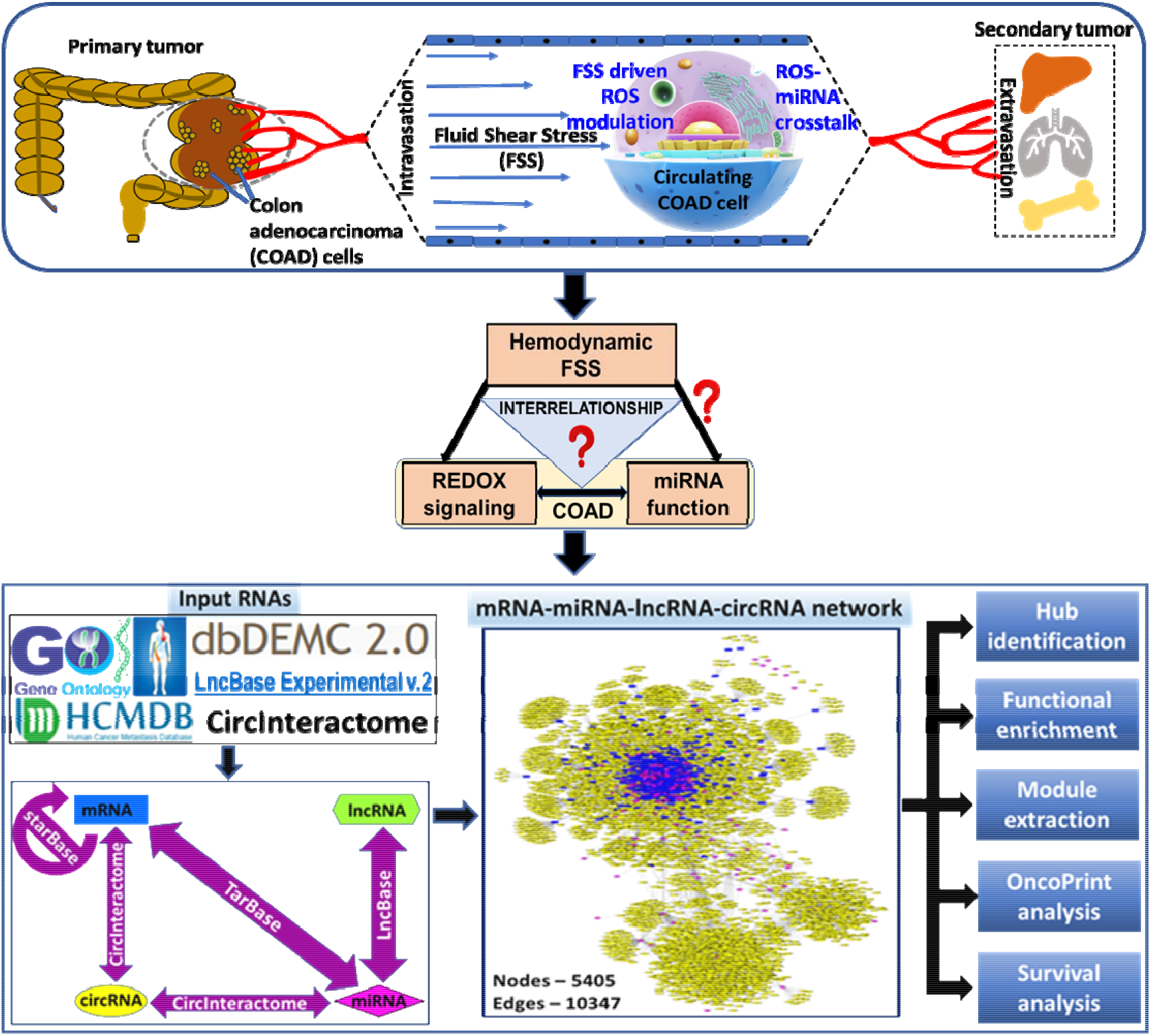
Outline of the integrated analysis.

**Figure 2.**
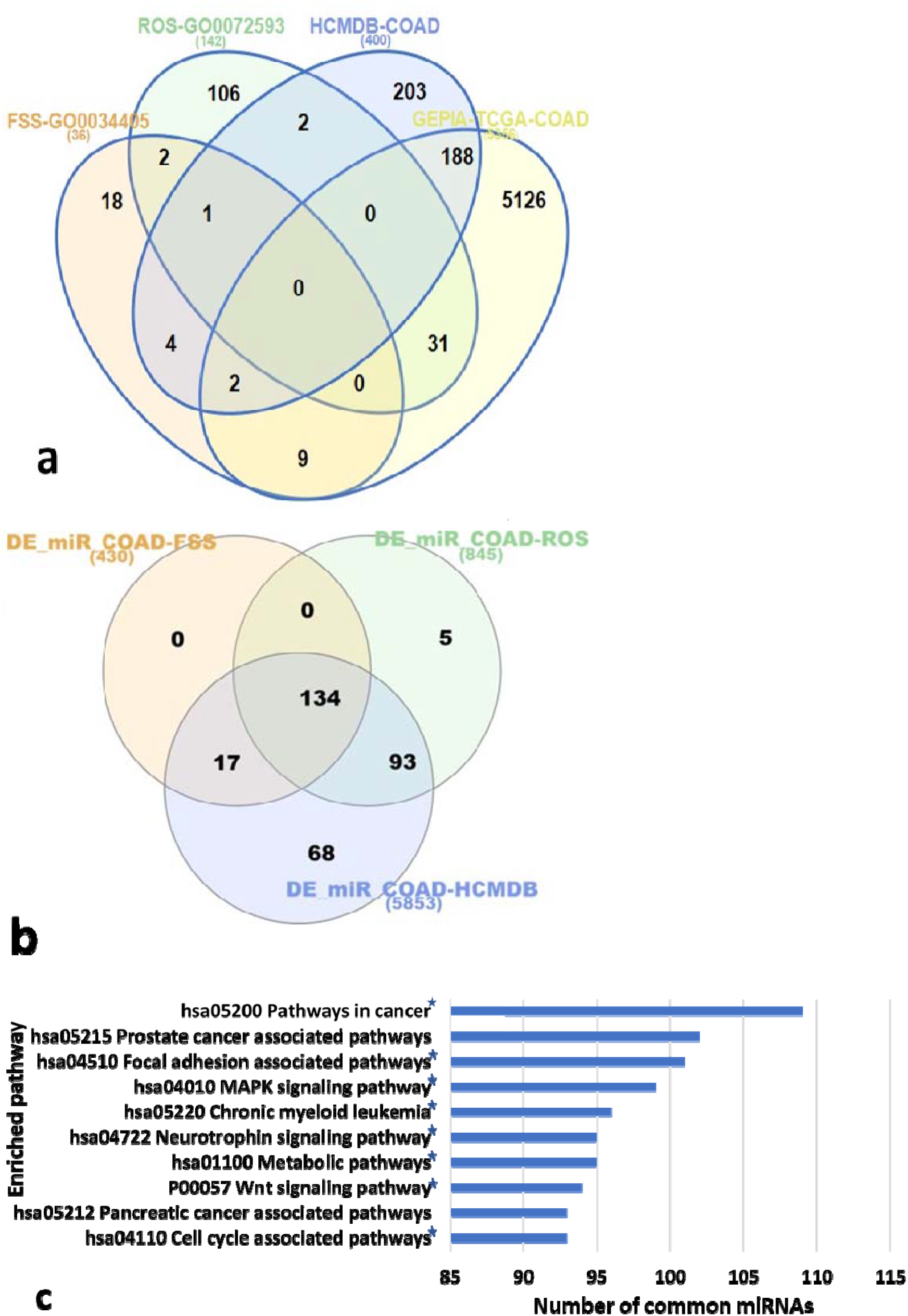
(a) Venn diagram representing all DE-COAD genes associated with FSS, ROS, HCMDB-COAD; (b) All the DE human miRNAs in COAD (DE_miR_COAD), interacting with the genes associated with FSS, ROS and HCMDB-COAD; and (c) Top 10 significant enriched pathways of 134 common miRNAs of FSS, ROS and HCMDB-COAD (p < 0.05). Here the Y-axis represents the number of common miRNAs (out of 134 identified) that the pathway is enriched with. * indicates that the pathway was reported to be involved in COAD metastasis based on the literature survey performed.

### Generation of interaction networks using all the selected RNAs and the expression network of DE-RNAs as well as RNA hub identification

Based on the interaction data retrieved, the mRNA-mRNA-miRNA networks for each gene list (FSS, ROS, and HCMDB-COAD) separately as well as for the all genes together (FSS+ROS+HCMDB-COAD (FRM) gene list) were generated and visualized using Cytoscape 3.7.1 [31] (Supplementary Figures 2a-2d). Interactions of lncRNAs and circRNAs were later merged to the above mentioned FRM network to generate a complex FRM mRNA-miRNA-lncRNA-circRNA network (Supplementary Figure 5). Finally, the top 20 RNA hubs, ranked based on their degree, of the networks were identified using cytoHubba app of Cytoscape 3.7.1 [43].

### Generation of a network of selected DE-RNAs in COAD, RNA hub identification and exploring their physiological relevance

DE mRNAs and DE miRNAs in COAD were identified using GEPIA [36] and dbDEMC 2.0 [35] respectively and the DE circRNAs in COAD were identified from GSE126094 dataset of Gene Expression Omnibus (GEO) database using GEO2R interactive web tool (with adjusted P-value < 0.05 and |log (2) FC|=2) [44]. The only lncRNA in this network, CCAT2, was reported to be upregulated in COAD [16, 45]. Based on this expression data of all the mRNAs, miRNAs, lncRNA and circRNAs, a more concise sub-network was generated (Supplementary Figure 6) and the top 20 degree-based RNA hubs of this expression network were also depicted using cytoHubba app [43]. Later, the biological significance of the identified RNA hubs was explored for their relevance in COAD metastasis using a literature survey.

### Identification, extraction and analyses of the functional modules of RNAs associated with FSS/ROS

The miRNAs associated with HCMDB-COAD (experimentally validated) [34] and the common DE-miRNAs between FSS, ROS, and HCMDB-COAD, enriched in GO terms “cellular response to fluid shear stress” and “reactive oxygen species metabolic process” obtained in miEAA [40] were added to the FRM mRNA-mRNA-miRNA and the FRM mRNA-miRNA-lncRNA-circRNA networks. Subsequently, module analysis of the resulting networks was performed using PEWCC1.0 plug-in of Cytoscape 3.7.1 [46].

### OncoPrint and survival analyses of the selected mRNAs and miRNAs

Since in network analysis, the significance of the molecules (nodes) in the process is determined by the number of interactions (i.e., degree), the biological importance was obtained through functional enrichment analysis, OncoPrint analysis and overall as well as metastasis-free survival analyses. Although the FSS and ROS associated mRNAs were obtained from GO, their alteration frequency in COAD patients (i.e., OncoPrint), as well as the correlation between the expression levels and survival of COAD patients, were investigated in our analysis for both mRNA and miRNAs. While the overall survival analysis of miRNAs associated with FSS, ROS, and HCMDB-COAD was done using starBase v2.0 [38], the survival analysis and the OncoPrint analysis of the mRNAs associated with the three phenomena were performed using PROGgeneV2 [47] and cBioPortal [48], respectively.

## Results and discussion

### Functional enrichment analysis of the selected miRNAs followed by an exploration of the biological significance of the miRNA-enriched pathways and the FSS/ROS associated mRNAs w.r.t. COAD metastasis

In our study, 190 HCMDB-COAD genes, 31 ROS genes, and 9 FSS genes that were DE in COAD were deduced from the Venn diagram analysis of mRNAs pertaining to FSS, ROS, HCMDB-COAD and GEPIA-DE-COAD exclusively (Figure 2a and Supplementary Table 1). Furthermore, the gene AKT1 was identified to be common among FSS, ROS, and COAD metastasis, but it was not found as DE mRNA in COAD. Also, four FSS associated genes (MTSS1, TGFβ1, SRC, PTGS2) and two ROS associated genes (EGFR and SESN2) emerged as involved in COAD metastasis. KLF4 and ASS1, the two DE FSS genes in COAD found in our study, were also reported to be associated with metastasis. When a literature survey of all the genes associated with either FSS or ROS or both was performed individually on Google Scholar and PubMed, 60% of the total 150 non-redundant genes were reported to play vital roles in COAD metastasis (Supplementary Table 2). This indicates that both FSS and ROS that we considered for our analysis are closely linked to COAD metastasis and also the bonding between them is highly significant. Likewise, 134 common miRNAs of the interacting COAD DE-miRNA partners of mRNAs associated with FSS, ROS, and HCMDB-COAD were identified (Figure 2b and Supplementary Table 3). When functional enrichment analysis of these 134 miRNAs was performed, the MAPK signaling pathway [3] and the Wnt signaling pathway [49] were found to be significant (Figure 2c). 9 out of the top 10 significant pathways, which were enriched with almost 70% or more of 134 common miRNAs, were also involved in COAD metastasis as evident from the existing literature (Supplementary Table 4). 99 common miRNAs were identified to be involved in the MAPK signaling pathway in our analysis, which increases cell migration and extravasation in MDA-MB-231 breast cancer cells via FSS-driven and ROS-induced activation of ERK1/2 of MAPK signaling pathway [10]. The Wnt signaling pathway, enriched with 94 common miRNAs that were identified, was another noteworthy biologically significant result of our analysis as FSS-mediated upregulation of β-catenin increased cell proliferation of HCT116 COAD cells [49]. The TGF-β signaling pathway that induced the CRC patient tumor cells to undergo metastasis via STAT3 signaling or activin signaling pathways [50, 51], showed up among the top 5 pathways under two different categories in our study (the Reactome pathways and GO) (Supplementary Figures 1a,1b) and was enriched with 83 COAD DE-miRNAs. Similarly, the p53 signaling pathway and cell migration were other noteworthy findings among the top 5 most significant Reactome pathways and GO, respectively (Supplementary Figures 1a,1b). Mutations in p53 have been reported in more than 60% of COAD cases and mutant p53 together with TGF-β suppression has been shown to promote intravasation leading to metastasis in a tumor mouse model [52]. Cell migration is a significant process that emerged in our study which involves the dynamics of actin polymerization and contraction that are inevitably required for the cells to undergo metastasis [53, 54] and also to acquire resistance against FSS [55] in the circulatory microenvironment. 83 and 72 common COAD DE-miRNAs linked to the p53 signaling pathway and cell migration process emerged in the study. The fact that the miRNA enriched pathways identified in this analysis, include not only those which suppress COAD metastasis (TGF-β and p53 signaling pathways) but also those which promote the process (MAPK signaling pathway, Wnt signaling pathway, cell migration) implies the biological relevance of the selected miRNAs obtained from the preliminary analysis of the study and substantiates them to be considered for further investigation. Altogether, 115 FSS/ROS associated miRNAs that play important roles in COAD metastasis were identified in our analysis (Supplementary Table 4).

**Table 1:**
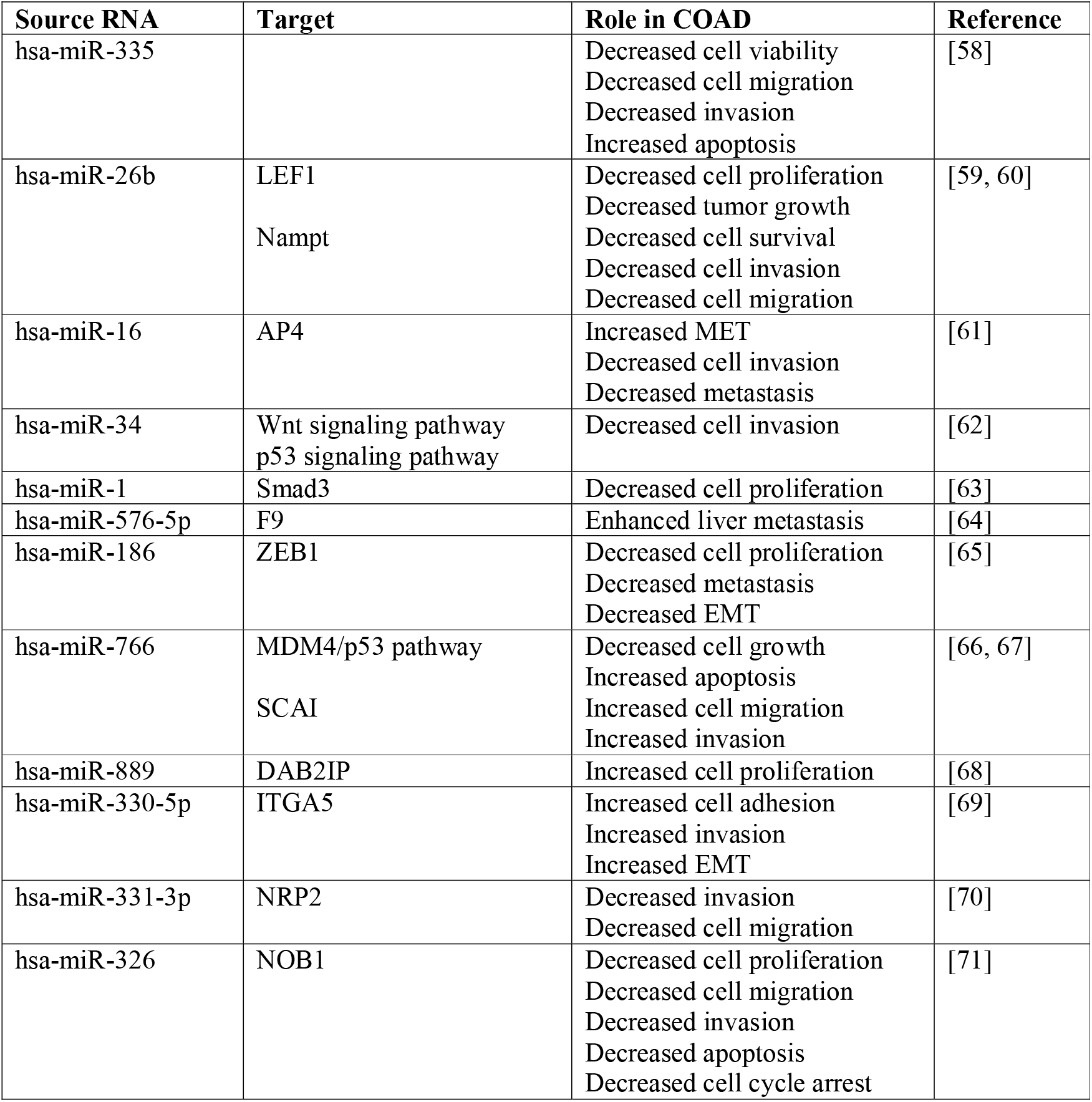
Validated target mRNAs and associated pathways of the selected miRNA hubs identified in the study with their role in COAD metastasis.

### Generation of various interaction networks of the selected RNAs, degree-based RNA hub identification and elucidation of their biological relevance to COAD metastasis

The interaction network comprised of 643 edges and 324 nodes was generated when the ROS associated mRNAs, as well as the COAD DE-miRNAs were considered (Supplementary Figure 2a). Similar networks associated with FSS showed 296 edges connecting 183 nodes (Supplementary Figure 2b), and that of HCMDB-COAD showed 4512 edges connecting 681 nodes, respectively (Supplementary Figure 2c). The merged network of these individual networks resulted in the FRM mRNA-mRNA-miRNA network with 798 nodes and 5280 edges (Supplementary Figure 2d); subsequently, the top 20 RNA hubs in each of the above four networks (mRNA-mRNA-miRNA networks corresponding to FSS, ROS, HCMDB-COAD, and FRM) were identified based on the degree (Supplementary Figures 3a, 3b, and 3c and Supplementary Tables 6, 7 and 8). Thus 10 miRNAs (hsa-miR-335-5p, hsa-miR-26b-5p, hsa-miR-16-5p, hsa-miR-34a-5p, hsa-miR-17-5p, hsa-miR-1-3p, hsa-miR-106b-5p, hsa-let-7b-5p, hsa-miR-92a-3p and hsa-miR-20a-5p) among the top 20 degree-based RNA hubs of the FRM network were identified (Figure 3a and Supplementary Table 9). When common RNAs between the top 20 RNA hubs of the four networks and the DE mRNAs in COAD were analyzed (Supplementary Figure 4, Supplementary Table 10), it showed that HMGA1, CCND1, RAN, HSP90AA1, and HNRNPA1 were the common RNAs between the top 20 HCMDB-COAD and the top 20 FRM networks. hsa-miR-26b-5p, a DE-miRNA in COAD, was identified to be common among all the four top 20 RNA hub networks, which suggests that this miRNA may play an important role in the process.

**Figure 3.**
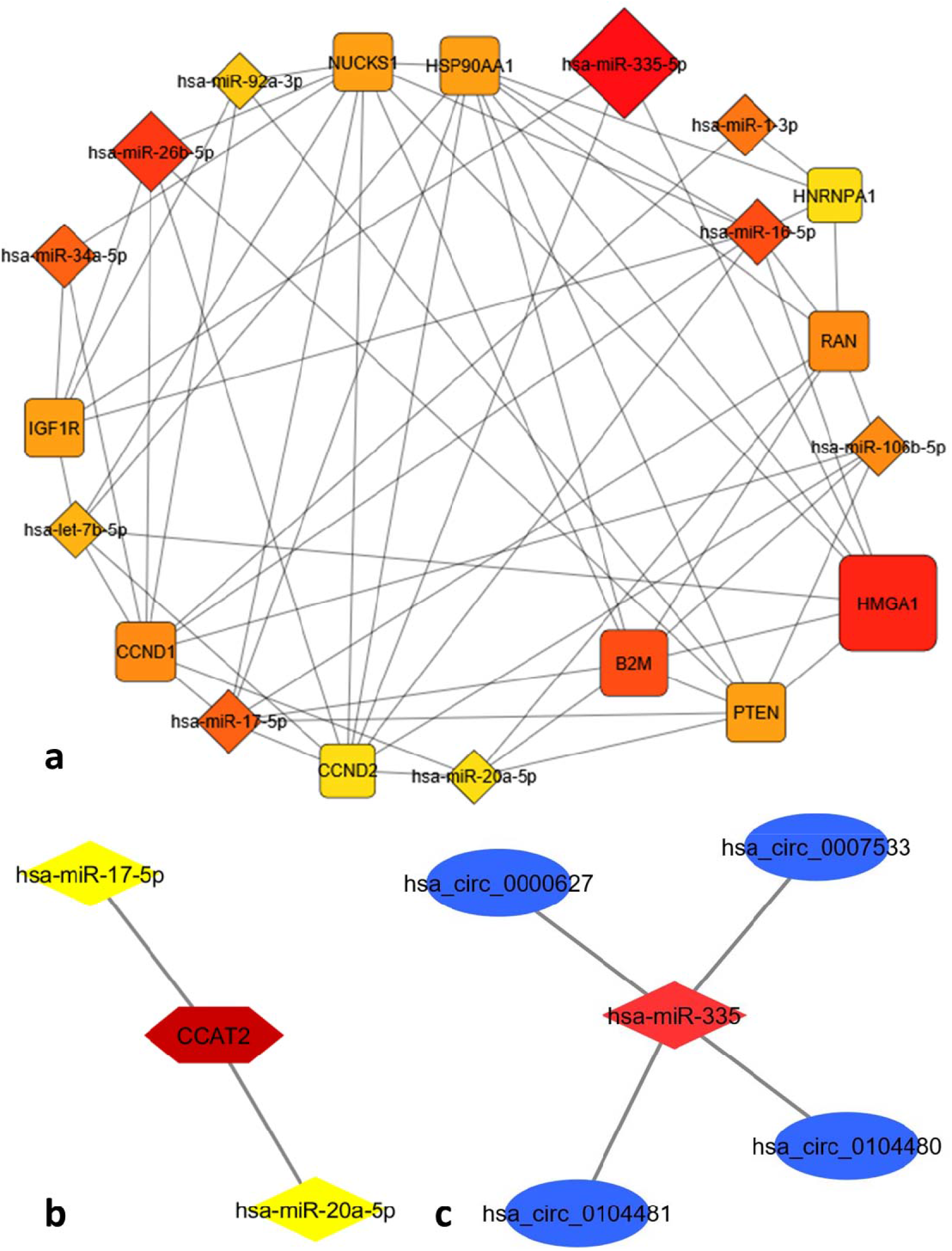
(a) Degree based top 20 miRNA and mRNA hubs of the merged FSS-ROS-HCMDB COAD mRNA-mRNA-miRNA network [The intensity gradient of the node color (ranging from dark orange denoting highest degree to yellow denoting lowest degree) of the top 20 hubs in the figure corresponds to the decreasing number of degree of the hub node]; (b) lncRNA-miRNA network [lncRNA: Brown hexagon; miRNA: Yellow diamond]; (c) circRNA-miRNA network [circRNA: Blue ellipse; miRNA: Red diamond]

When significant lncRNAs and circular RNAs that interact with the miRNA hubs of the FRM mRNA-mRNA-miRNA network were explored, a single lncRNA (CCAT2) was identified to interact with hsa-miR-17-5p, and hsa-miR-20a-5p (Figure 3b) and four circular RNAs (hsa_circ_0000627, hsa_circ_0007533, hsa_circ_0104480, and hsa_circ_0104481) were identified to interact with hsa-miR-335 (Figure 3c). This implies that CCAT2, which is a downstream target in the Wnt signaling, highly expressed in COAD patient samples and showed a positive correlation with metastatic progression and a negative correlation with the overall survival rates [16, 45, 56], showed interactions with two significant miRNA hubs of the FRM network. Though the physiological role of recently discovered circRNAs is still unexplored, elucidation of the circRNAs that were identified in the present study as associated with FSS-driven and ROS-mediated COAD metastasis, particularly those that showed differential expression in COAD, may provide novel therapeutic strategies.

FRM mRNA-miRNA-lncRNA-circRNA network showed 10347 edges connecting 5405 nodes (481 mRNAs, 351 miRNAs, 1 lncRNA and 4572 circular RNAs) (Supplementary Figure 5) and degree-based top 20 RNA hubs of this network include nine miRNAs (hsa-miR-186, hsa-miR-766, hsa-miR-576-5p, hsa-miR-335-5p, hsa-miR-889, hsa-miR-331-3p, hsa-miR-330-5p, hsa-miR-326 and hsa-miR-26b-5p) (Figure 4a and Supplementary Table 11). The emergence of all these miRNAs that regulate various steps of COAD metastasis as topmost miRNA hubs in our network analysis exhibits the robustness of these results, details of which were described in Table 1. The expression network consisted of 2059 interactions between 518 DE RNAs in COAD (208 DE mRNAs, 291 DE miRNAs, 1 DE lncRNA and 18 DE circRNAs) (Supplementary Figure 6). Among the mRNAs of the expression network within the top 20 RNA hubs (Figure 4b), BCL2, which modulates COAD progression via the Wnt signaling pathway, emerged as a noteworthy mRNA associated with ROS [49]. The pro-apoptotic gene PMAIP1 (NOXA) that increases the sensitivity of COAD cells to chemotherapy [57] was another ROS associated mRNA identified in our study. Glycogen synthase kinase-3β (GSK-3β), that gets suppressed by the FSS-driven ROS/NO production and promote EMT and metastasis in breast cancer cells [9] showed up as one of the most significant mRNA hubs in the expression network. Five miRNAs (hsa-miR-335-5p, hsa-miR-16-5p, hsa-miR-26b-5p, hsa-miR-34a-5p, hsa-miR-1-3p) were identified in the top 20 degree-based RNA hubs of the expression network (Figure 4b and Supplementary Table 12). Among all the miRNA hubs identified, hsa-miR-335 was observed as the most significant one with the highest degree in most of the networks of this study.

**Figure 4.**
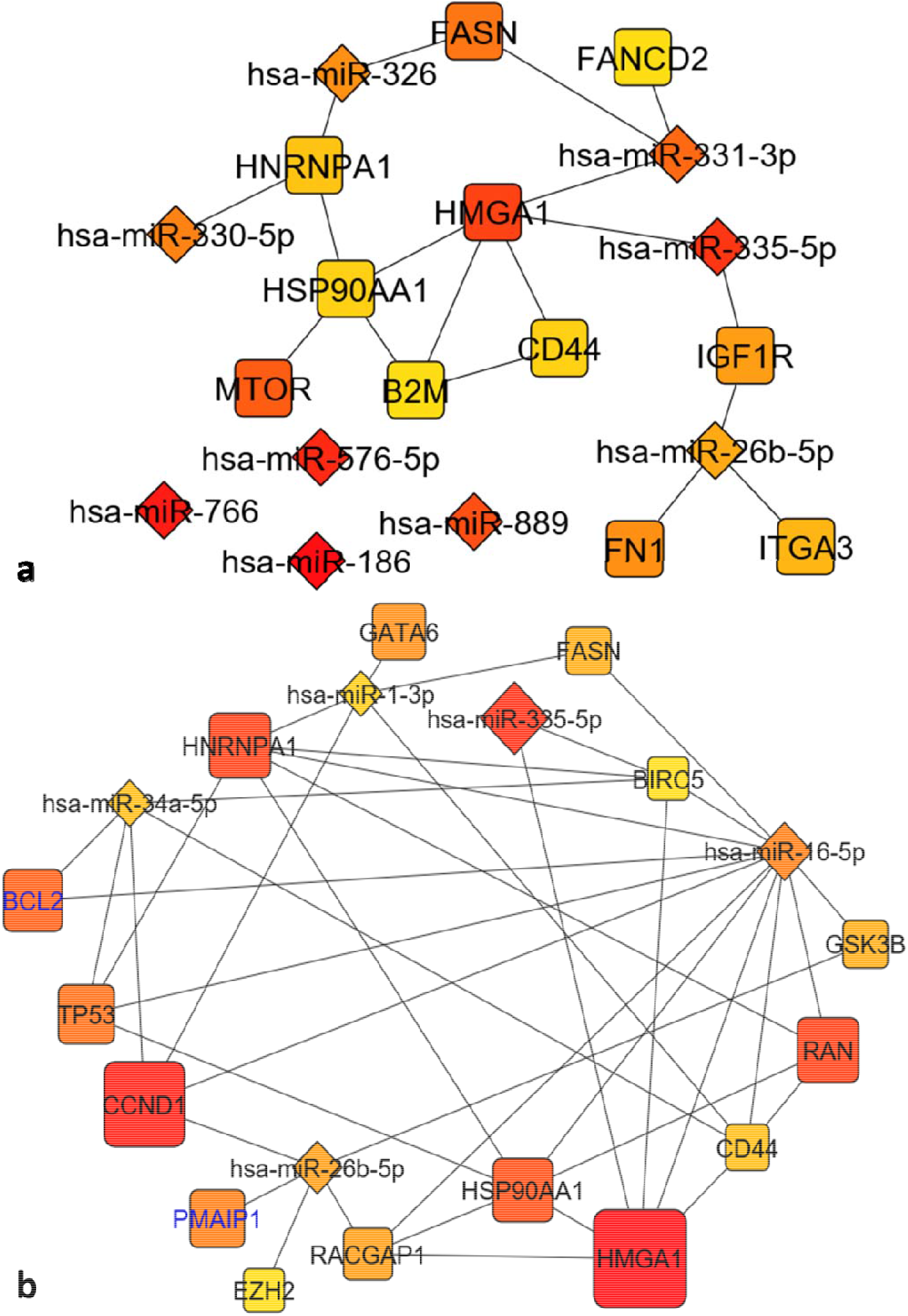
(a) Identification of degree-based top 20 RNA hubs of the FRM COAD mRNA-miRNA-lncRNA-circRNA network (b) Identification of degree-based top 20 RNA hubs of FRM COAD mRNA-miRNA-lncRNA-circRNA expression network; [Node size (largest to smallest) is proportional to the degree (highest to lowest); ROS mRNA nodes in blue font color]. In figures 4 (a-b) – The intensity gradient of the node color (ranging from dark orange denoting highest degree to yellow denoting lowest degree) of the top 20 hubs in the figure corresponds to the decreasing number of degree of the hub node.

### Functional module analyses and identification of the mRNAs/miRNAs that link FSS, ROS and HCMDB-COAD

Although the generated networks provide the comprehensive view of all the interactions, small subnetworks (modules) that comprise only the most significant RNA hubs associated with various phenomena are essential to perform the validation experiments [46]. Hence, module analysis was performed based on the concept of weighted cluster coefficient [46] to see whether the miRNAs identified in the study were directly associated with FSS and/or ROS or not (based on the GO data obtained during miEAA [40] analysis as shown in Supplementary Table 5). When module analysis was performed, 575 modules that included RNAs directly associated with at least two of the three phenomena (FSS, ROS and HCMDB-COAD) were identified and extracted from the FRM mRNA-mRNA-miRNA network. Among them, six FSS/ROS associated modules (Figure 5a-5f), two ROS/HCMDB-COAD associated modules (Supplementary Figures 7a, 7b), and two FSS/HCMDB-COAD associated modules were also identified (Supplementary Figures 7c, 7d). In these modules, while four miRNAs (hsa-miR-185-5p, hsa-miR-21-5p, hsa-miR-335-5p and hsa-miR-145-5p) were associated with both ROS and HCMDB-COAD, six (hsa-miR-451a, hsa-miR-150-5p, hsa-miR125b-5p, hsa-miR-143-3p, hsa-miR-186-5p and hsa-miR-335-5p) were pertaining to FSS and ROS. Similarly, four miRNAs (hsa-miR-375, hsa-miR-335-5p, hsa-miR-26b-5p and hsa-miR-143-3p) showed an association with FSS and HCMDB-COAD. hsa-miR-335-5p, hsa-miR-143-3p, and hsa-miR-150-5p that were observed as miRNA hubs in more than one sub-network may act as novel molecular markers in FSS and ROS mediated COAD metastasis.

**Figure 5 (a-f).**
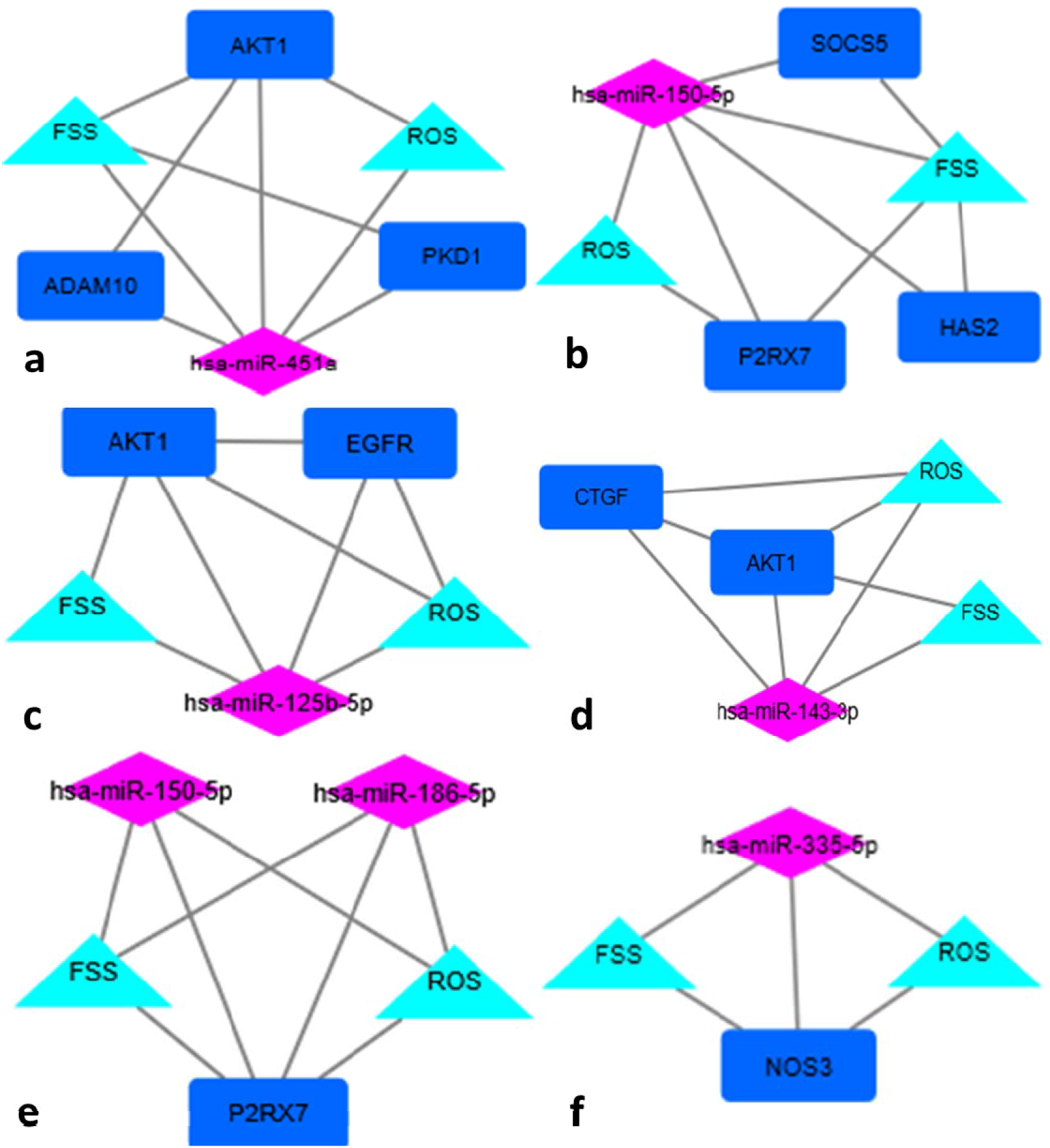
Module analysis in FRM mRNA-mRNA-miRNA network [pink diamond-miRNA; dark blue-mRNA; Aquablue-phenomenon (FSS/ROS)]

### OncoPrint analysis and survival analyses of the selected mRNAs/miRNAs pertaining to FSS and ROS

The OncoPrint analysis of the FSS and ROS associated mRNAs that was performed using 594 COAD patient sample data from TCGA [48] revealed 15 mRNAs with more than 10% gene alterations in 80% of the patients/samples (SRC-31%, IMMP2L-15%, MTSS1-15%, PXDNL-15%, PREX1-15%, EGFR-14%, LRRK2-14%, ADAM9-13%, ABCA1-12%, HAS2-12%, SLC7A2-12%, GBF1-12%, PXDN-12%, POR-11%, AKT1-11%) (Figure 6a). Subsequently, the OncoPrint analysis performed using 1134 Metastasis-COAD patient sample data showed nine mRNAs that exhibited variable types of gene alterations in 16% of the patients/samples (SRC-5%, EGFR-4%, PDGFRB-3%, AKT1-1.6%, BCL2-1.5%, FANCC-1.3%, PMAIP1-1.1%, KLF4-1.1%, and NFE2L2-0.8%) (Supplementary Figure 8). Among these, the most significant mRNAs were SRC, EGFR, and AKT1 as they showed gene alterations in both TCGA as well as the metastasis-COAD patient samples. SRC [72] and EGFR [73], which played crucial roles in COAD progression, are associated with FSS/HCMDB-COAD and ROS/HCMDB-COAD respectively. Furthermore, the fact that AKT1 [74], that regulates COAD metastasis, altered in both TCGA-COAD as well as COAD-metastasis and showed association with FSS, ROS, and HCMDB-COAD is one of the most prominent results. Another noteworthy result of the OncoPrint analysis was the emergence of the transporter proteins, ABCA1 and SLC7A2 (Figure 6a), which mediated cell invasion and tumor progression in COAD cells respectively [75, 76]. Our result that SLC7A2 showed significant gene alteration in COAD but not in metastasis is supported by the literature [76]. Thus, the significant gene alterations of the FSS and ROS associated mRNAs observed in COAD patients indicate that these mRNAs may be used as prognostic biomarkers of COAD metastasis.

**Figure 6.**
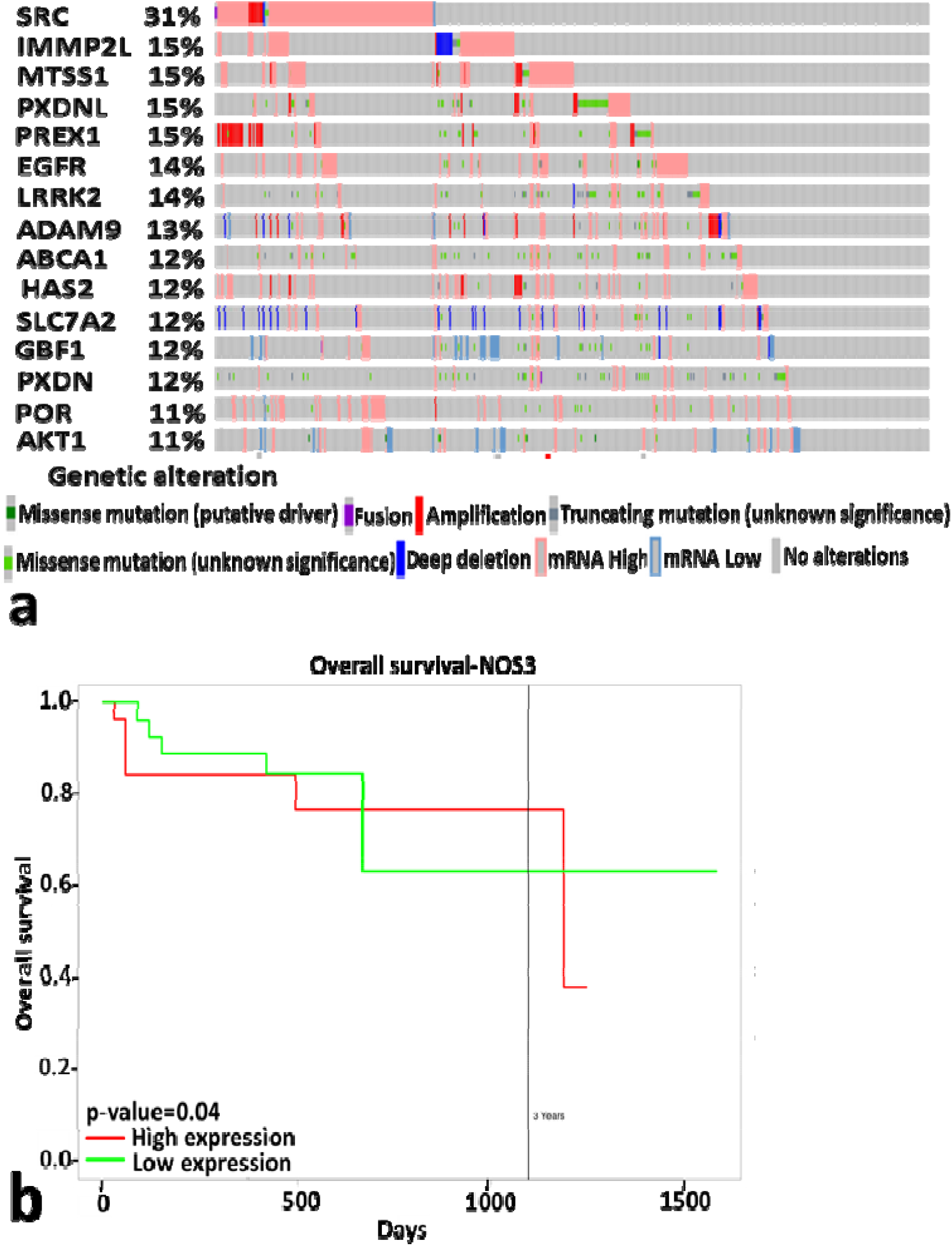
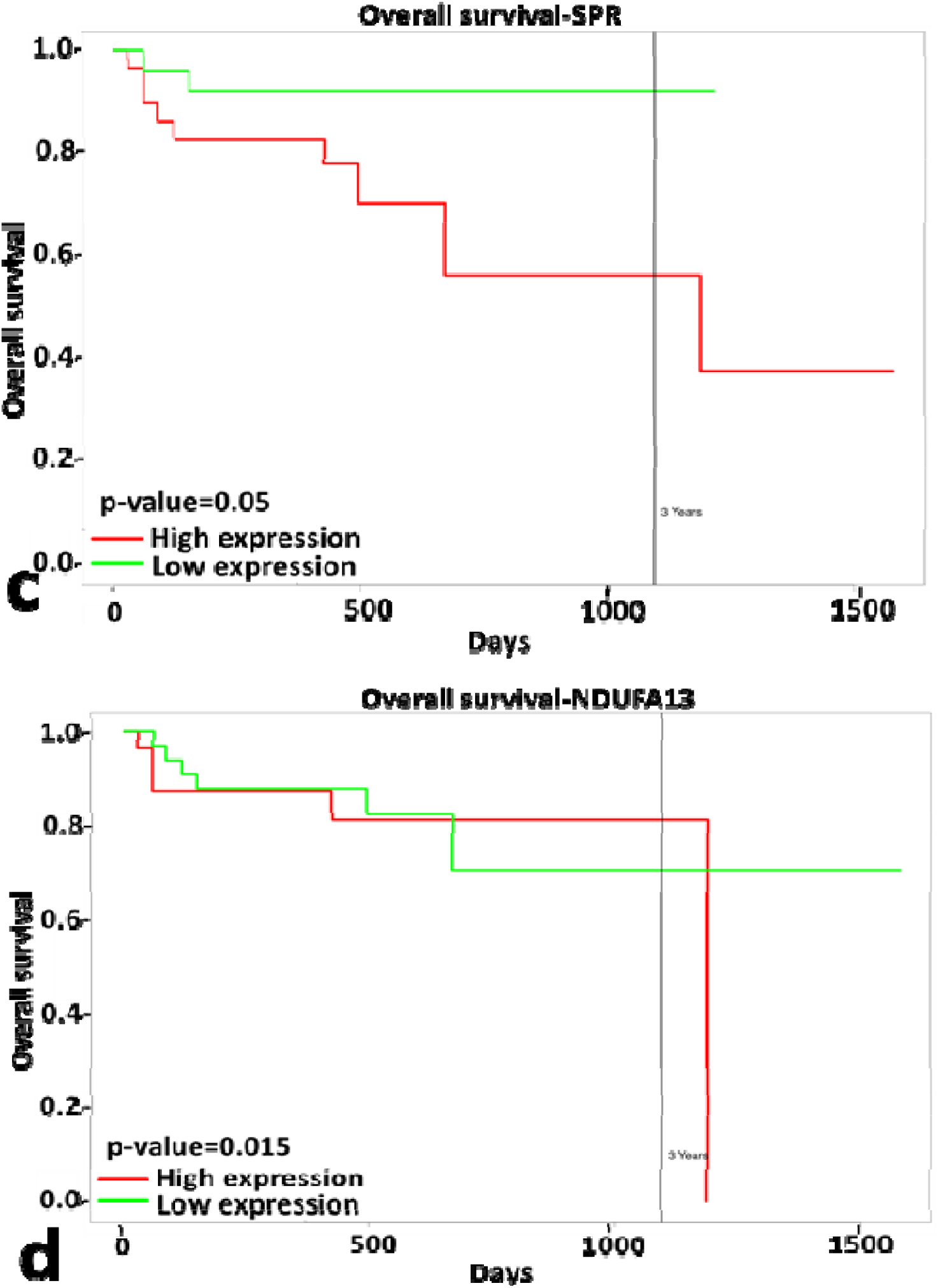
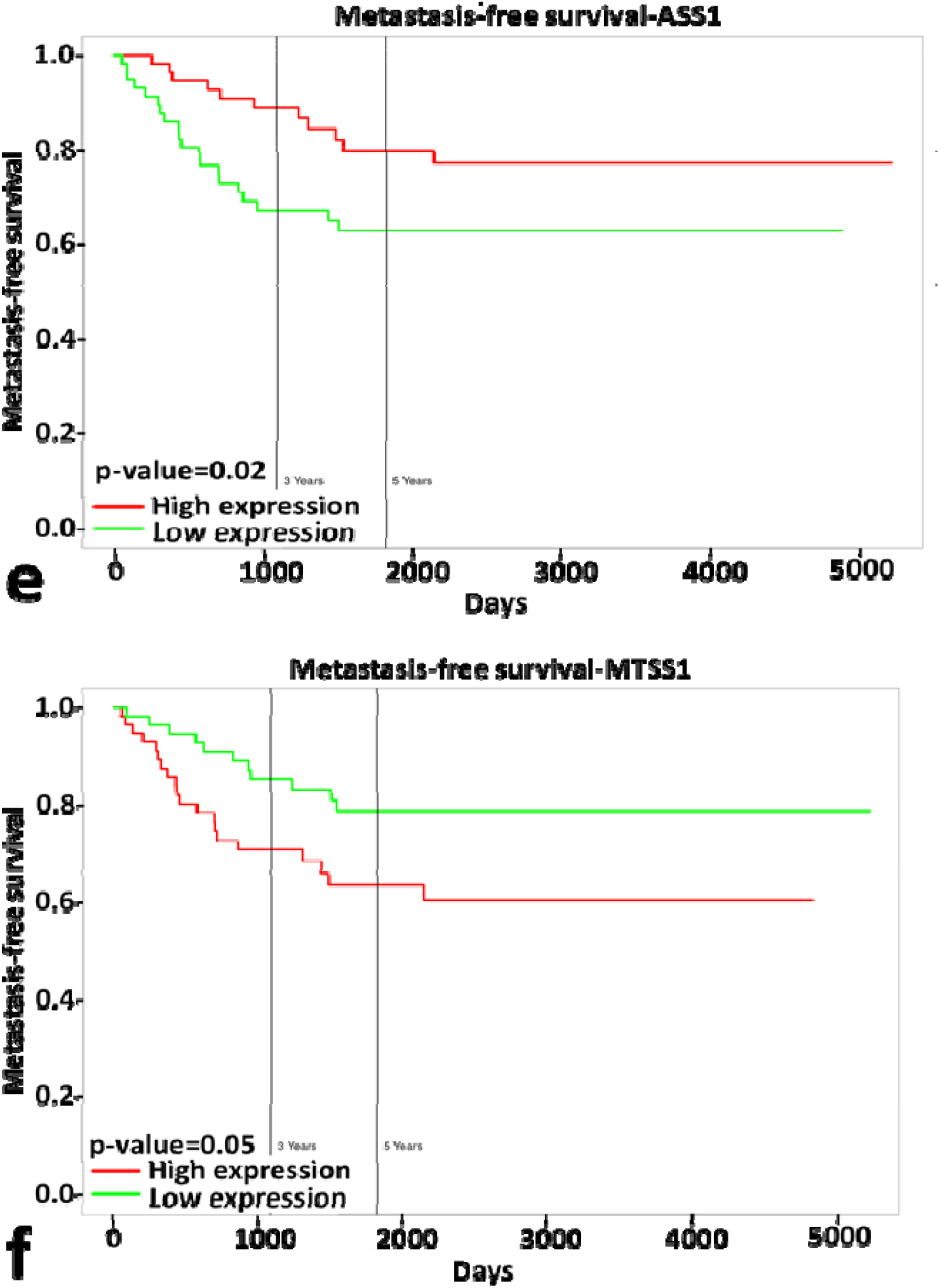
OncoPrint and survival analyses of significant mRNAs associated with FSS, ROS, and HCMDB-COAD. OncoPrint analyses of 15 significant mRNAs obtained using TCGA data (Fig. 6a); overall and metastasis-free survival plots of 3 significant mRNAs (Fig. 6b-6d) and two significant mRNAs, respectively (Fig. 6e-6f)

When the overall survival analysis of the 134 common miRNA interacting partners of FSS, ROS and HCMDB-COAD was performed using TCGA sample data, 16 significant miRNAs (hsa-miR-200b-3p, hsa-miR-193b-3p, hsa-miR-130a-3p, hsa-miR-92a-3p, hsa-miR-497-5p, hsa-miR-708-5p, hsa-miR-503-5p, hsa-miR-552-3p, hsa-miR-181b-5p, hsa-miR-375, hsa-miR-193a-3p, hsa-miR-200a-5p, hsa-miR-200c-5p, hsa-miR-21-3p, hsa-miR-210-3p and hsa-miR-497-3p with log-rank p < 0.05) in COAD were identified. However, the most significant miRNAs among them which also showed a direct association with either both FSS and ROS (hsa-miR-193b-3p and hsa-miR-375) (Figures 7a and 7b) or with both ROS and HCMDB-COAD (hsa-miR-200b-3p) (Figure 7c) or with ROS (hsa-miR-181b-5p) (Figure 7d) based on miEAA result data (Supplementary Table 5), were identified as potential prognostic biomarkers in the process. When the overall survival analysis of the selected mRNAs in the TCGA COAD samples was done, NOS3 (both FSS and ROS associated) (Figure 6b), SPR (Figure 6c) (ROS-associated), and NDUFA13 (Figure 6d) (ROS-associated) were identified as significant (P < 0.05) overall survival markers. Analysis of the metastasis-free survival plots of the selected mRNAs revealed 33 significant metastasis-free survival markers in COAD (with log-rank p < 0.05) based on GSE28814/GSE28722 data. Out of these, 22 ROS-associated mRNAs (NOS2, GPX3, MAOB, GCHFR, GCH1, HBB, PDK4, PXDN, RFK, NOX4, CCS, DDAH2, PDGFB, EDN1, CTGF, PRDX2, SPR, PAX2, PLA2R1, CYP1A1, NDUS1, PRDX3), 7 FSS-associated mRNAs (TGFB3, MEF2C, PKD2, SMAD6, SOCS5, CSF2, PDGFRB), two mRNAs associated with both FSS and HCMDB-COAD (ASS1, MTSS1) (Figures 6e and 6f) were identified as the most significant. The association of the expression profiles of all these miRNAs and mRNAs with the metastasis-free survival (Supplementary Table 13) and overall survival (Supplementary Table 14) status in TCGA sample data were also enlisted separately and could be used as survival biomarkers in COAD metastasis.

**Figure 7.**
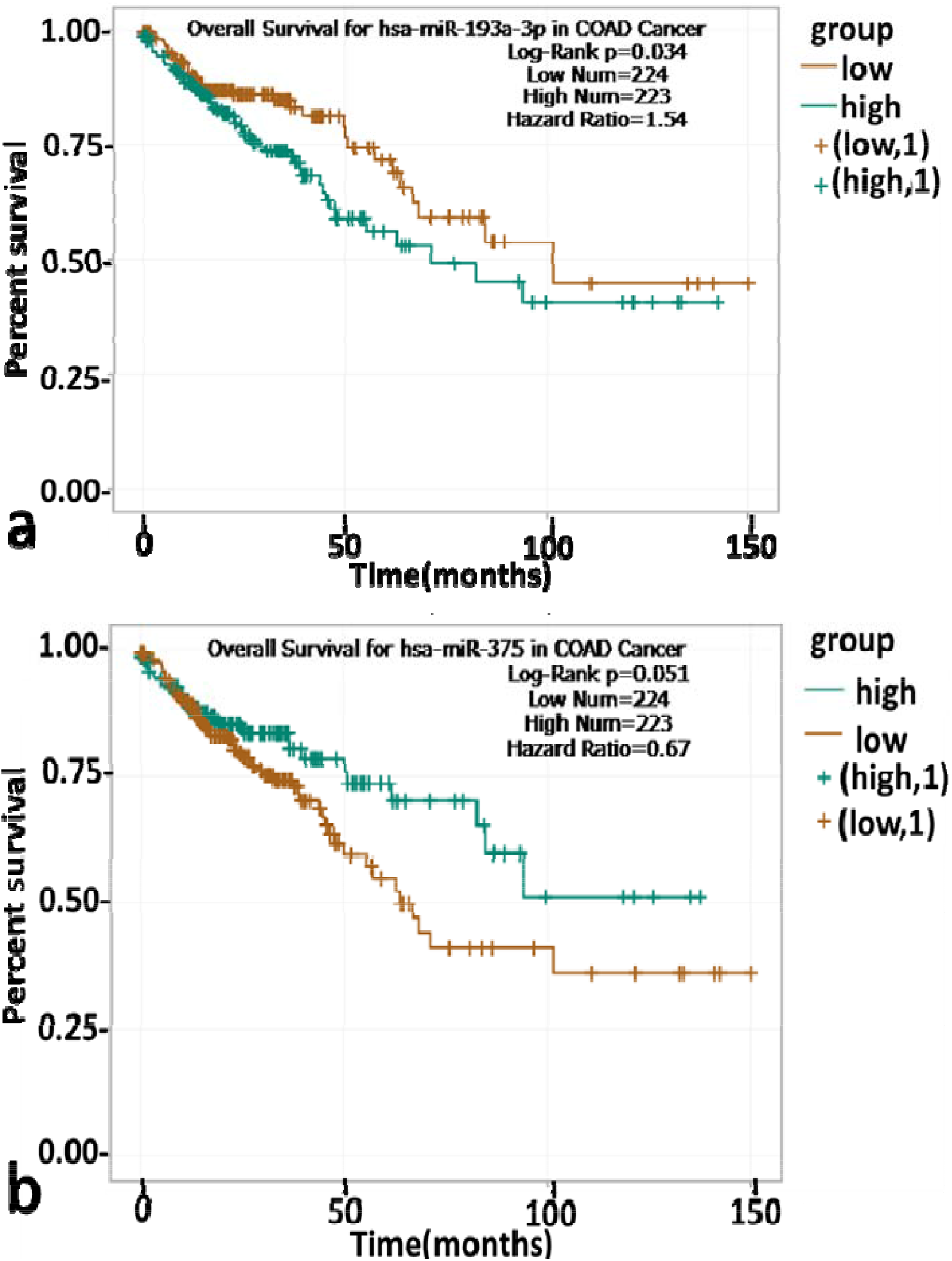
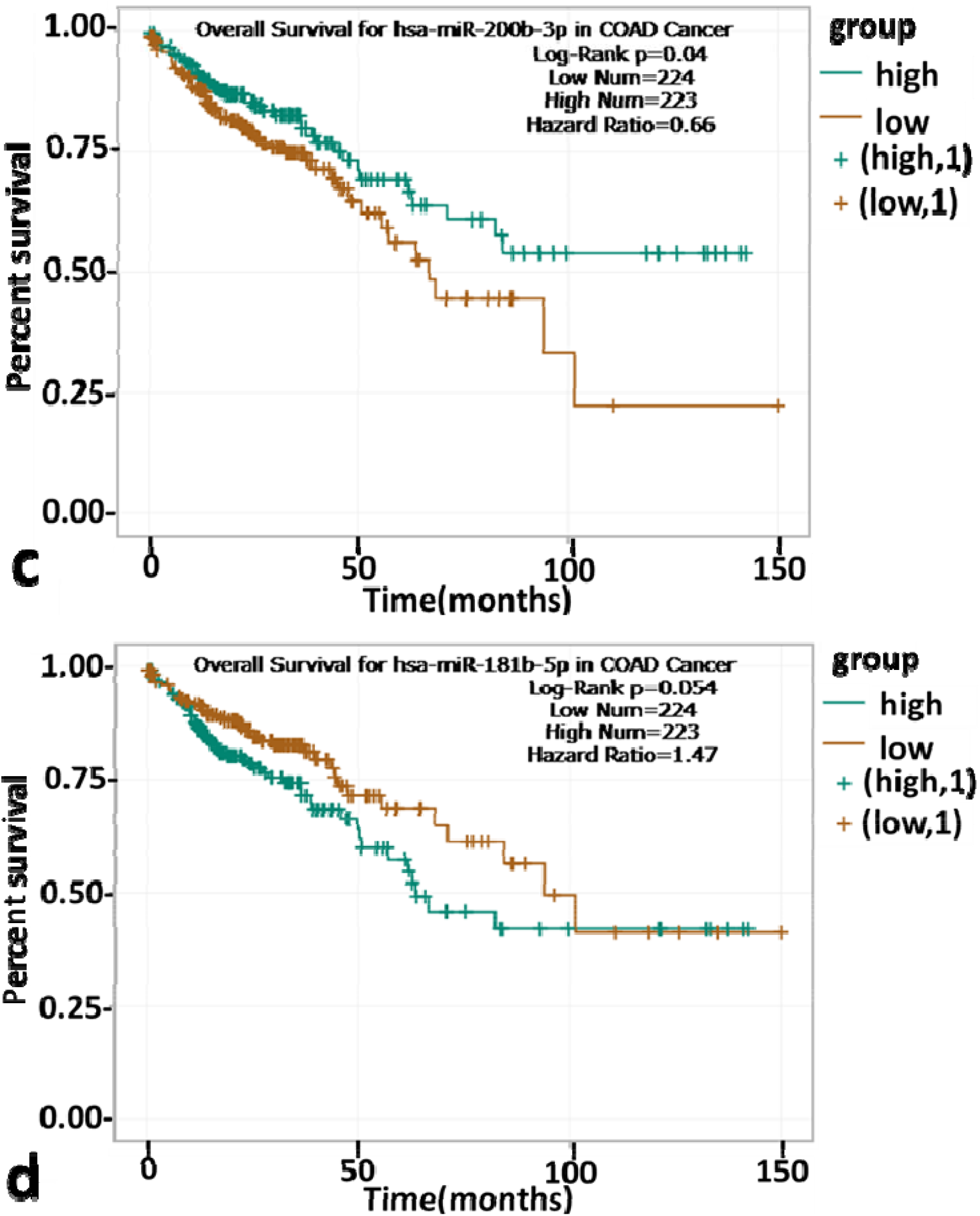
Overall survival plots of 4 significant miRNAs associated with FSS and/or ROS in COAD (Fig. 6a-d)

### Clinical significance of FSS, ROS and miRNAs in other cancers with relevance to our study in COAD metastasis

The synergistic effect of FSS and ROS enhances apoptosis of the circulatory tumor cells of the lung, breast, and cervical cancer when exposed to FSS and treated with ROS-generating anticancer drugs such as doxorubicin or cisplatin simultaneously [12]. Elevated superoxide dismutase (SOD) levels in the circulatory breast cancer cells mediated molecular changes that reduced FSS-driven and superoxide mediated apoptosis and increased resistance against the anti-cancer drug, doxorubicin [77]. In addition to the known roles in apoptosis, DNA repair, cell cycle regulation, and cancer stem cell formation, certain miRNAs also act as key regulators in the process of acquiring multidrug resistance in COAD cells [78]. Furthermore, the huge potential of the miRNAs to reduce cancer progression could be exploited to design miRNA-targeted therapeutics using mimics of tumor suppressor miRNAs or anti-mimics of the oncomiRNAs [79, 80]. The mimics of hsa-miR-34 (MRX34), a tumor suppressor and a potential therapeutic agent for COAD has entered the phase I clinical trials also [80]; miR-34a-5p was found as one of the most significant hubs in our expression network analysis.

Thus, the present study unveiled several physiologically and clinically significant coding/non-coding RNAs involved in FSS-driven and ROS-mediated COAD metastasis. Further experimental validation of the interactions among the most significant mRNAs (such as SRC, EGFR and AKT1), miRNAs (such as hsa-miR-16-5p, hsa-miR-335-5p, hsa-miR-26b-5p, hsa-miR-34a-5p and hsa-miR-1-3p), lncRNA (CCAT2) and the DE circRNAs identified in the study may provide highly valuable insights to explore novel strategies for efficient cancer treatment and management.

## Conclusion

In our study, FSS, ROS, and non-coding RNAs were studied together to obtain a broad overview of the molecular interactions that may occur in the circulating microenvironment associated with COAD metastasis. To achieve this, a novel approach in generating the interaction network involving various biological phenomena associated with the process was considered to understand the mechanism and was biologically correlated. Though the effect of FSS on the miRNAs of endothelial cells is extensively studied, the association between FSS and miRNAs in cancer pathogenesis is still unexplored. This study was performed to unmask the relationship between FSS and miRNAs in COAD cells and reveal the significance of this relationship in COAD metastasis and survival.

hsa-miR-16-5p, hsa-miR-335-5p, hsa-miR-26b-5p, hsa-miR-34a-5p, and hsa-miR-1-3p that were highlighted in the study as the significant miRNAs associated with the FSS, ROS, and HCMDB-COAD, also showed differential expression in COAD. The lncRNA, CCAT2, and the circular RNAs, hsa_circ_0000627, hsa_circ_0007533, hsa_circ_0104480 and hsa_circ_0104481 that were reported in this study, also may play significant regulatory roles in the FSS-driven and ROS-mediated COAD metastasis.

Module analysis performed to obtain selected subnetworks comprising significant RNA hubs directly associated with FSS, ROS, and HCMDB-COAD not only paved the way to understand the molecular interactions better but also resulted in the identification of hsa-miR-335-5p and hsa-miR-150-5p as the most significant miRNA hubs involved in the process. Further, the OncoPrint analysis and the survival analyses (overall as well as metastasis-free survival) of the identified miRNAs and mRNAs, was also performed to study their importance in the process. The results showed that significant gene alterations of these RNAs occurred in the COAD patients and that the expression status of the same also influenced the survival of the patients, thus emphasizing on their clinical significance as well.

The outcome of this study that includes the novel identification of hsa-miR-335-5p and hsa-miR-150-5p as molecular markers and hsa-miR-181b-5p, hsa-miR-193b-3p, hsa-miR-375 and hsa-miR-200b-3p as potential prognostic survival biomarkers, seems remarkable in FSS driven, and ROS mediated COAD metastasis. SRC, EGFR, and AKT1 were noteworthy among the mRNAs that showed significant gene alterations in COAD patients. Further experimental validation studies on these physiologically and clinically significant RNAs identified in our study may provide us valuable insights to design novel treatment and management strategies to reduce FSS driven and ROS mediated COAD metastasis-related deaths.

## Acknowledgements

The authors would like to thank Dr. Swagatika Sahoo, Department of Chemical Engineering, IIT Madras, India and Dr. Karthik Raman, Department of Biotechnology, Bhupat and Jyoti Mehta School of Biosciences, IIT Madras, India for providing invaluable inputs to fine-tune and strengthen the analysis.

## Author Contributions

SK and SB conceptualized the work, processed the data and performed the analyses. DK and GK oversaw the work and provided directions to improve it. All the authors worked on the manuscript together.

## Additional Information

### List of Supplementary figures

**<u>Supplementary figure 1a</u>**: Top 5 most significant Reactome pathways enriched with common miRNA interacting partners of FSS, ROS and HCMDB-COAD (p < 0.05)

**<u>Supplementary figure 1b</u>**: Top 5 most significant Gene Ontology terms enriched with common miRNA interacting partners of FSS, ROS and HCMDB-COAD (p < 0.05).

**<u>Supplementary figure 2a</u>**: ROS mRNA-mRNA-miRNA network

**<u>Supplementary figure 2b</u>**: FSS mRNA-mRNA-miRNA network

**<u>Supplementary figure 2c</u>**: HCMDB-COAD mRNA-mRNA-miRNA network

**<u>Supplementary figure 2d</u>**: Merged FRM mRNA-mRNA-miRNA network

**<u>Supplementary figure 3a</u>**: Degree based top 20 RNA hubs of ROS mRNA-mRNA-miRNA network

**<u>Supplementary figure 3b</u>**: Degree based top 20 RNA hubs of FSS mRNA-mRNA-miRNA network

**<u>Supplementary figure 3c</u>**: Degree based top 20 RNA hubs of HCMDB-COAD mRNA-mRNA-miRNA network

**<u>Supplementary figure 4</u>**: InteractiVenn diagram representing mRNA from degree-based top 20 hubs of individual FSS, ROS, HCMDB-COAD, merged FRM mRNA-mRNA-miRNA networks and the DE genes in COAD (GEPIA-TCGA-COAD)

**<u>Supplementary figure 5</u>**: FRM COAD mRNA-miRNA-lncRNA-circRNA network

**<u>Supplementary figure 6</u>**: FRM COAD DE mRNA-DE miRNA-DE lncRNA-DE circRNA expression network

**<u>Supplementary figure 7</u>**: Module analysis using FRM mRNA-mRNA-miRNA network as the input: Extraction of modules showing direct association of identified miRNAs with either ROS and HCMDB-COAD or FSS and HCMDB-COAD

**<u>Supplementary figure 8</u>**: OncoPrint analyses of nine significant mRNAs associated with FSS, ROS, and HCMDB-COAD obtained using MSKCC metastasis data.

### List of Supplementary tables

***<u>Supplementary Table 1:</u>*** Common and unique genes obtained from the InteractiVenn diagram representing genes associated with FSS, ROS, HCMDB-COAD and the DE genes in COAD (GEPIA-TCGA-COAD)

***<u>Supplementary Table 2:</u>*** Role of FSS and ROS associated genes in COAD metastasis based on a literature survey

***<u>Supplementary Table 3:</u>*** List of 134 DE miRNAs (DE-miRs) – common among [DE miR COAD-FSS] and [DE miR COAD-ROS] and [DE miR COAD-HCMDB COAD]

***<u>Supplementary Table 4:</u>*** Common miRNA-enriched pathways and their role in COAD metastasis based on a literature survey

***<u>Supplementary Table 5:</u>*** List of miRNAs directly associated with FSS and ROS based on miEAA result data

***<u>Supplementary Table 6:</u>*** Degree-based top 20 RNA hubs of ROS mRNA-mRNA-miRNA network

***<u>Supplementary Table 7:</u>*** Degree-based top 20 RNA hubs of FSS mRNA-mRNA-miRNA network

***<u>Supplementary Table 8:</u>*** Degree-based top 20 RNA hubs of HCMDB-COAD mRNA-mRNA-miRNA network

***<u>Supplementary Table 9:</u>*** Degree-based top 20 RNA hubs of merged FRM mRNA-mRNA-miRNA network

***<u>Supplementary Table 10:</u>*** Common mRNAs among top 20 degree-based hubs of individual FSS, ROS, HCMDB-COAD, FRM mRNA-mRNA-miRNA networks and the DE genes in COAD (GEPIA-TCGA-COAD)

***<u>Supplementary Table 11:</u>*** Degree-based top 20 RNA hubs of FRM COAD mRNA-miRNA-lncRNA-circRNA network

***<u>Supplementary Table 12:</u>*** Degree-based top 20 RNA hubs of FRM COAD DE mRNA-DE miRNA-DE lncRNA-DE circRNA network

***<u>Supplementary Table 13:</u>*** Variation of metastasis-free survival status of COAD patients with the expression levels of significant mRNAs associated with FSS and ROS

***<u>Supplementary Table 14:</u>*** Variation of survival status of COAD patients with the expression levels of significant miRNAs and mRNAs associated with FSS and ROS using TCGA data

