## Supplementary Figures for "Non-coding RNAs in fluid shear stress-driven and reactive oxygen species-mediated colon cancer metastasis"

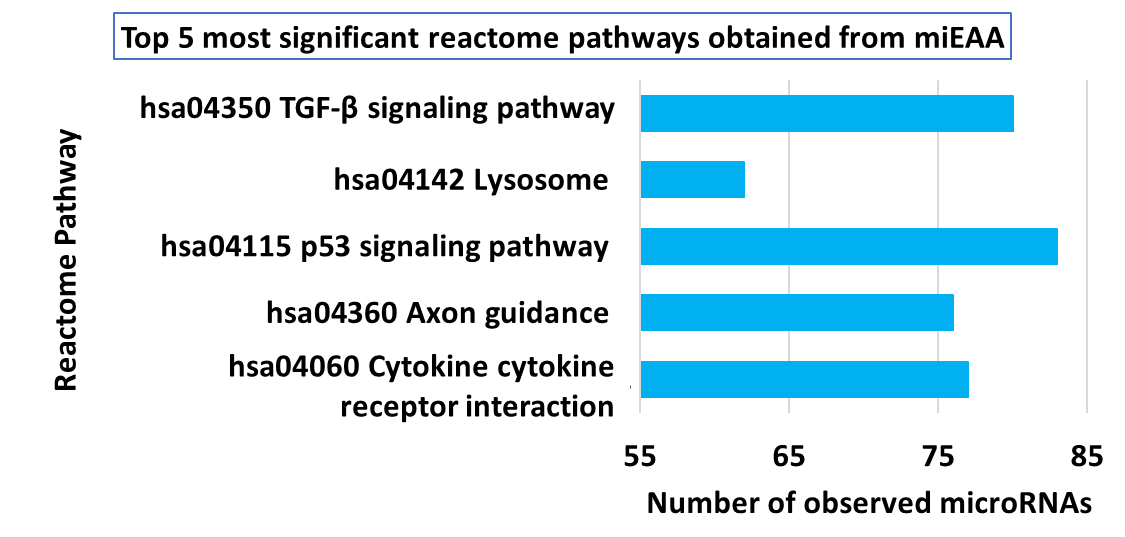

**Supplementary figure 1a: Top five most significant Reactome pathways enriched with common miRNA interacting partners of FSS, ROS and HCMDB-COAD (p < 0.05).**

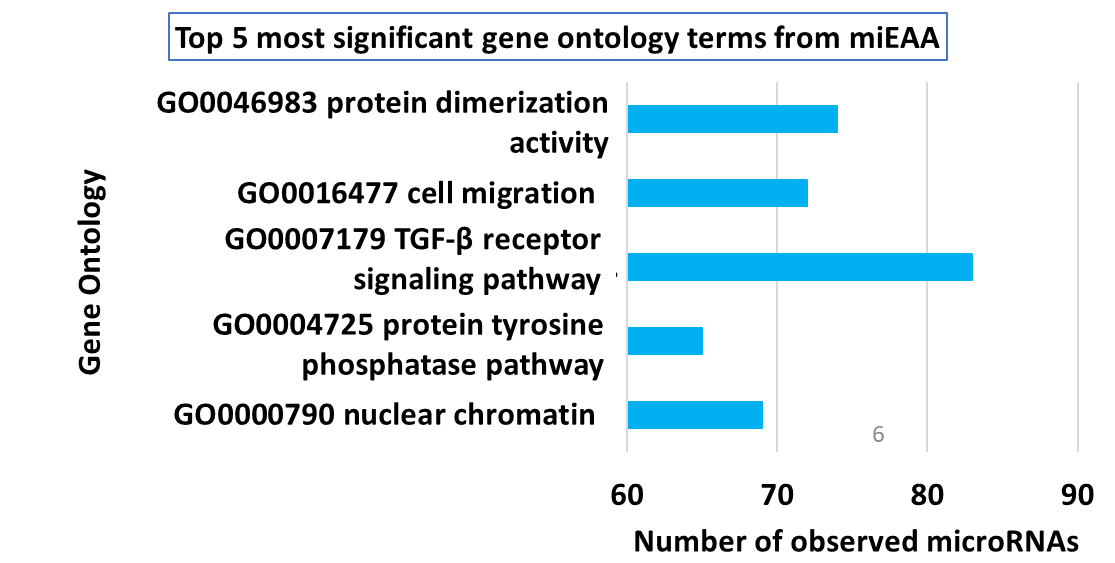

**Supplementary figure 1b: Top five most significant Gene Ontology terms enriched with common miRNA interacting partners of FSS, ROS and HCMDB-COAD (p < 0.05).**

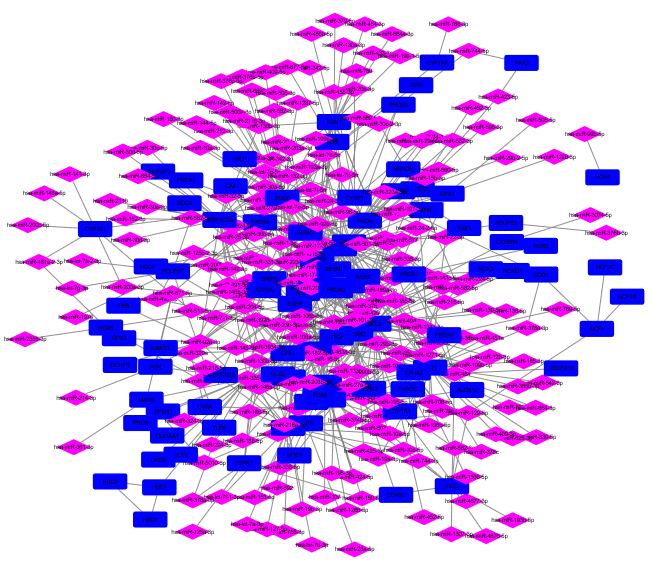

**Supplementary figure 2a: ROS mRNA-mRNA-miRNA network [mRNA and miRNA associated with ROS were indicated as dark blue rectangle and pink diamond respectively]**

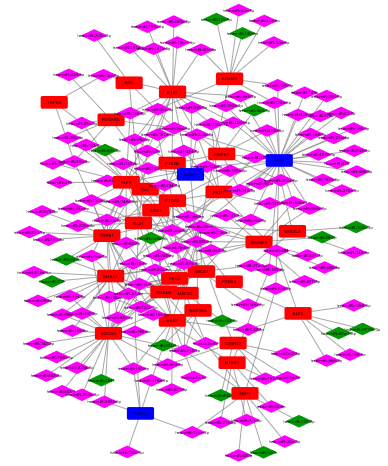

**Supplementary figure 2b: FSS mRNA-mRNA-miRNA network [mRNA and miRNA were indicated as rectangle and diamond respectively; red/dark green node colour – FSS; dark blue/pink node colour – common between FSS & ROS]**

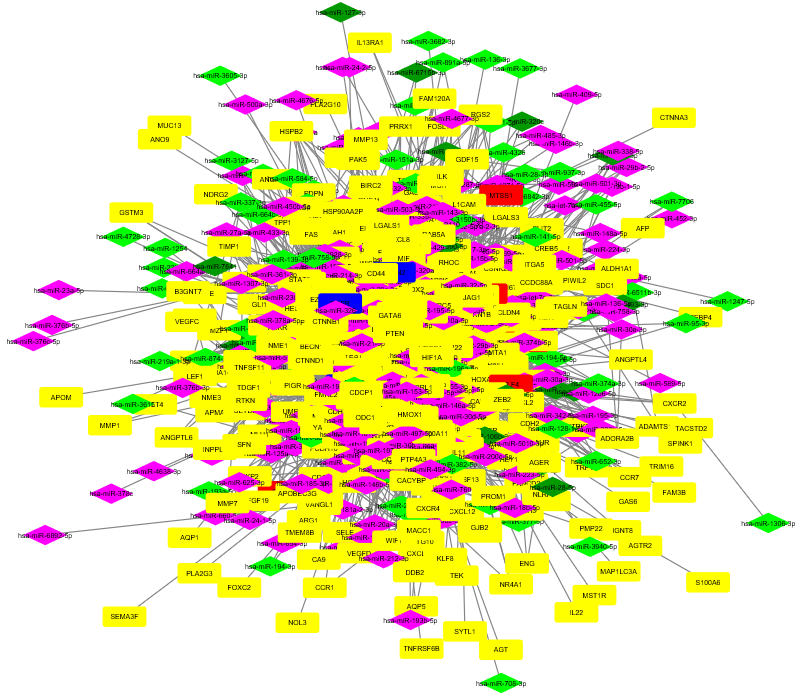

**Supplementary figure 2c: HCMDB COAD mRNA-mRNA-miRNA network [mRNA and miRNA were indicated as rectangle and diamond respectively; yellow/fluorescent green – unique to HCMDB COAD; dark blue/pink – common between HCMDB – COAD & ROS; red/dark green – common between HCMDB-COAD & FSS]**

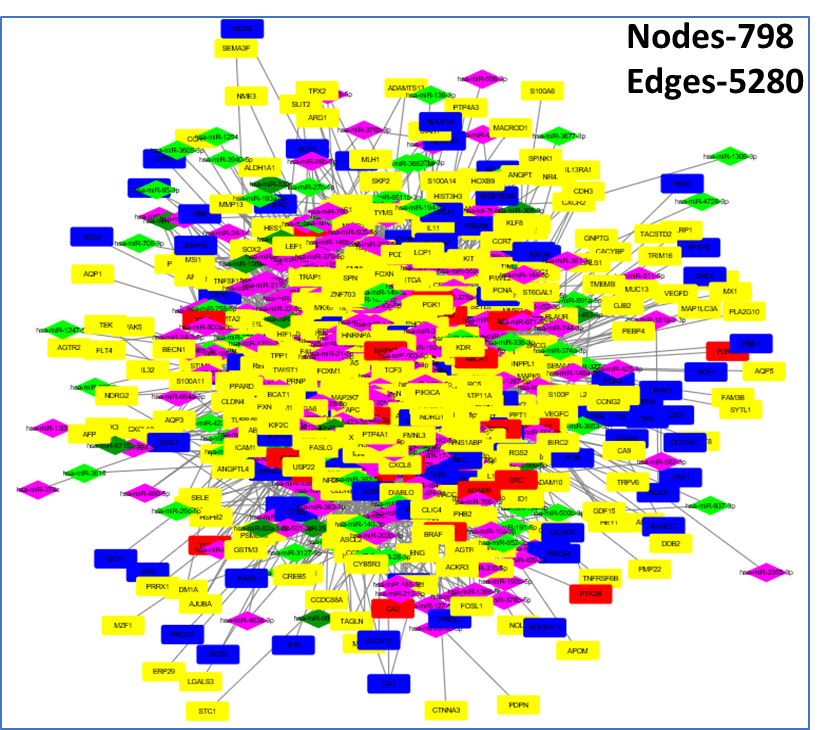

**Supplementary figure 2d: Merged FRM mRNA-mRNA-miRNA network [mRNA and miRNA were indicated as rectangle and diamond respectively; dark blue/pink – ROS; red/dark green – FSS; yellow/fluorescent green – HCMDB COAD]** **[FRM – FSS, ROS and HCMDB-COAD]**

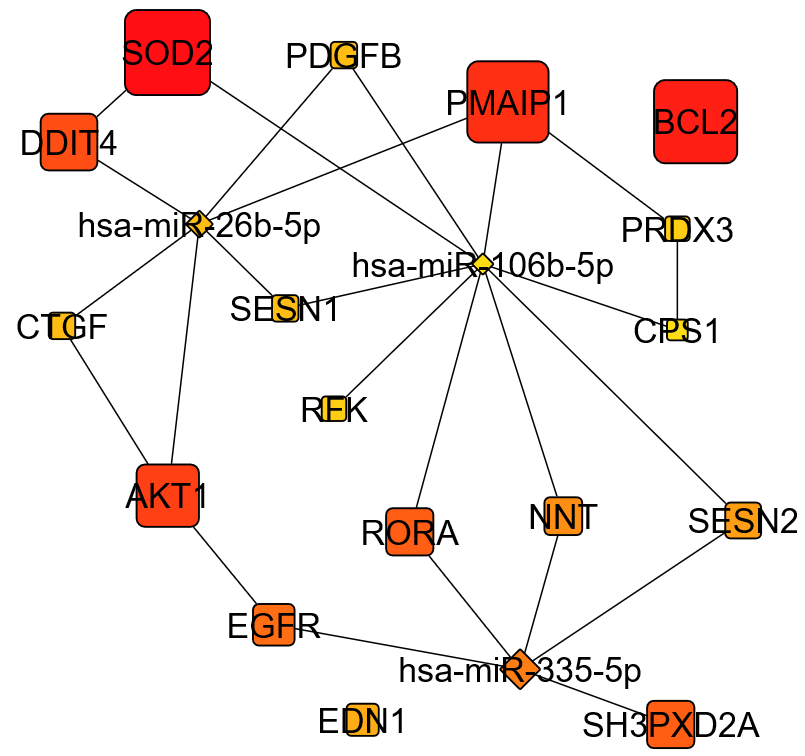

**Supplementary figure 3a: Degree based top 20 RNA hubs of** **ROS mRNA-mRNA-miRNA network** **[node size (largest to smallest) and node color intensity (dark orange to yellow) are proportional to the degree (highest to lowest)]**

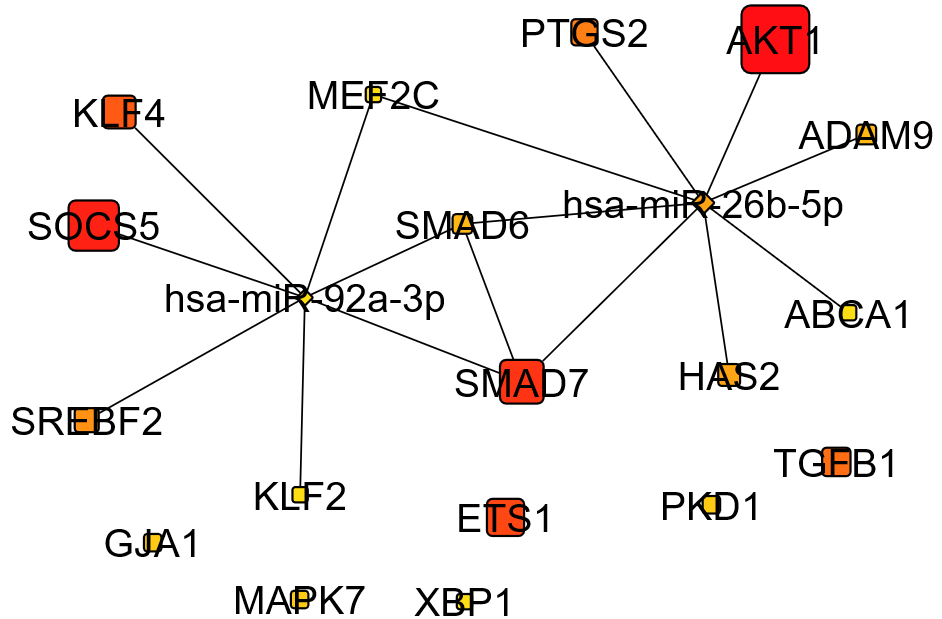

**Supplementary figure 3b: Degree based top 20 RNA hubs of** **FSS mRNA-mRNA-miRNA network** **[node size (largest to smallest) and node color intensity (dark orange to yellow) are proportional to the degree (highest to lowest)]**

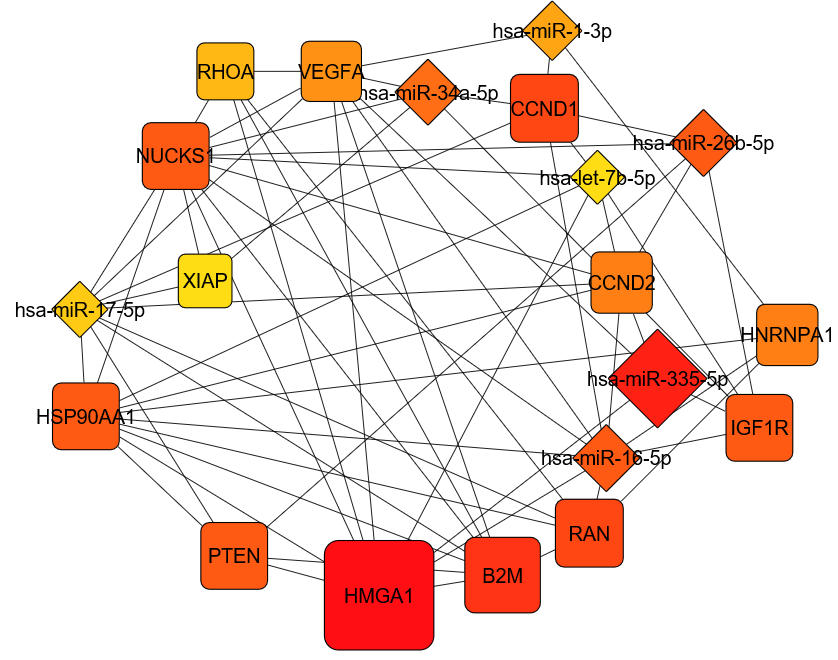

**Supplementary figure 3c: Degree based top 20** **RNA hubs of** **HCMDB-COAD mRNA-mRNA-miRNA network** **[node size (largest to smallest) and node color intensity (dark orange to yellow) are proportional to the degree (highest to lowest)]**

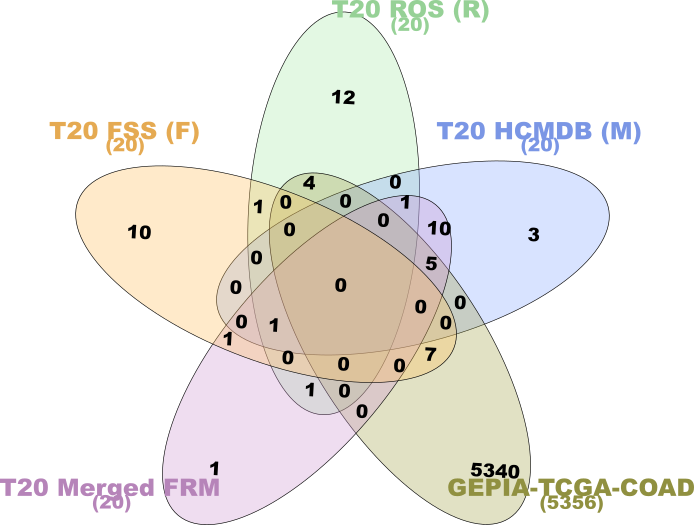

**Supplementary figure 4: InteractiVenn diagram representing mRNA from degree based top 20 hubs of individual FSS, ROS, HCMDB-COAD, merged FRM mRNA-mRNA-miRNA networks and the DE genes in COAD (GEPIA-TCGA-COAD) [FRM – FSS, ROS and HCMDB-COAD]**

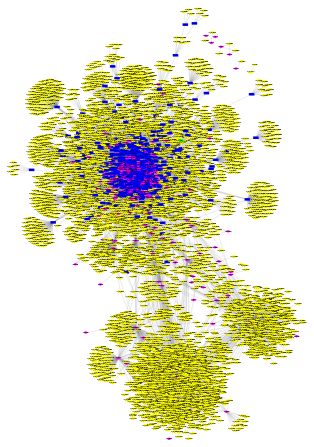

**Supplementary figure 5: FRM COAD mRNA-miRNA-lncRNA-circRNA network** **[circRNA: yellow ellipse; lncRNA: fluorescent green hexagon; mRNA: dark blue rectangle; miRNA: pink diamond] [FRM – FSS, ROS and HCMDB-COAD]**

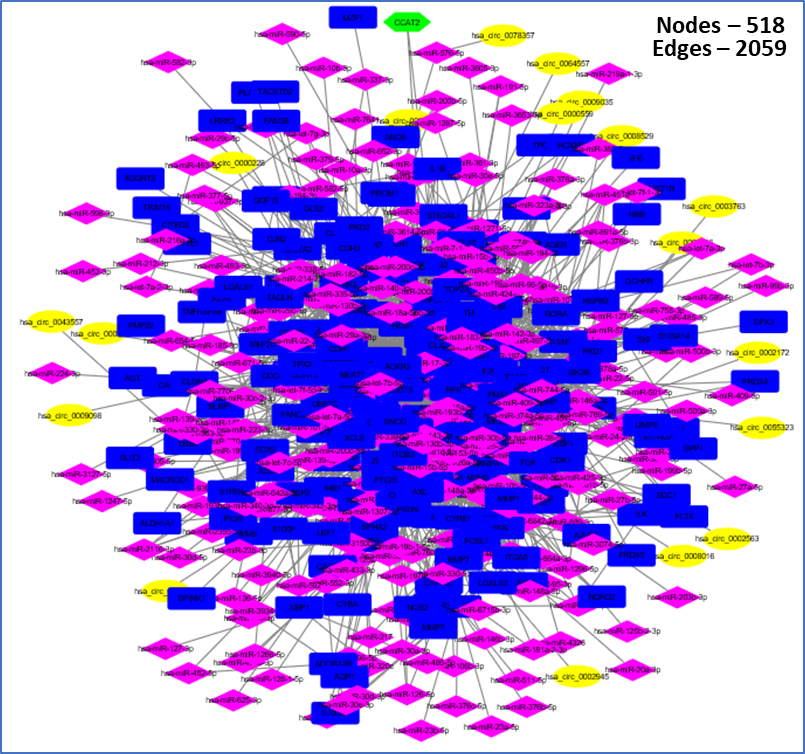

**Supplementary figure 6: FRM COAD DE mRNA-DE miRNA-DE lncRNA-DE circRNA expression network [DE mRNA: blue rectangle; DE miRNA: pink diamond; DE lncRNA: fluorescent green hexagon; DE circRNA: yellow ellipse] [FRM – FSS, ROS and HCMDB-COAD]**

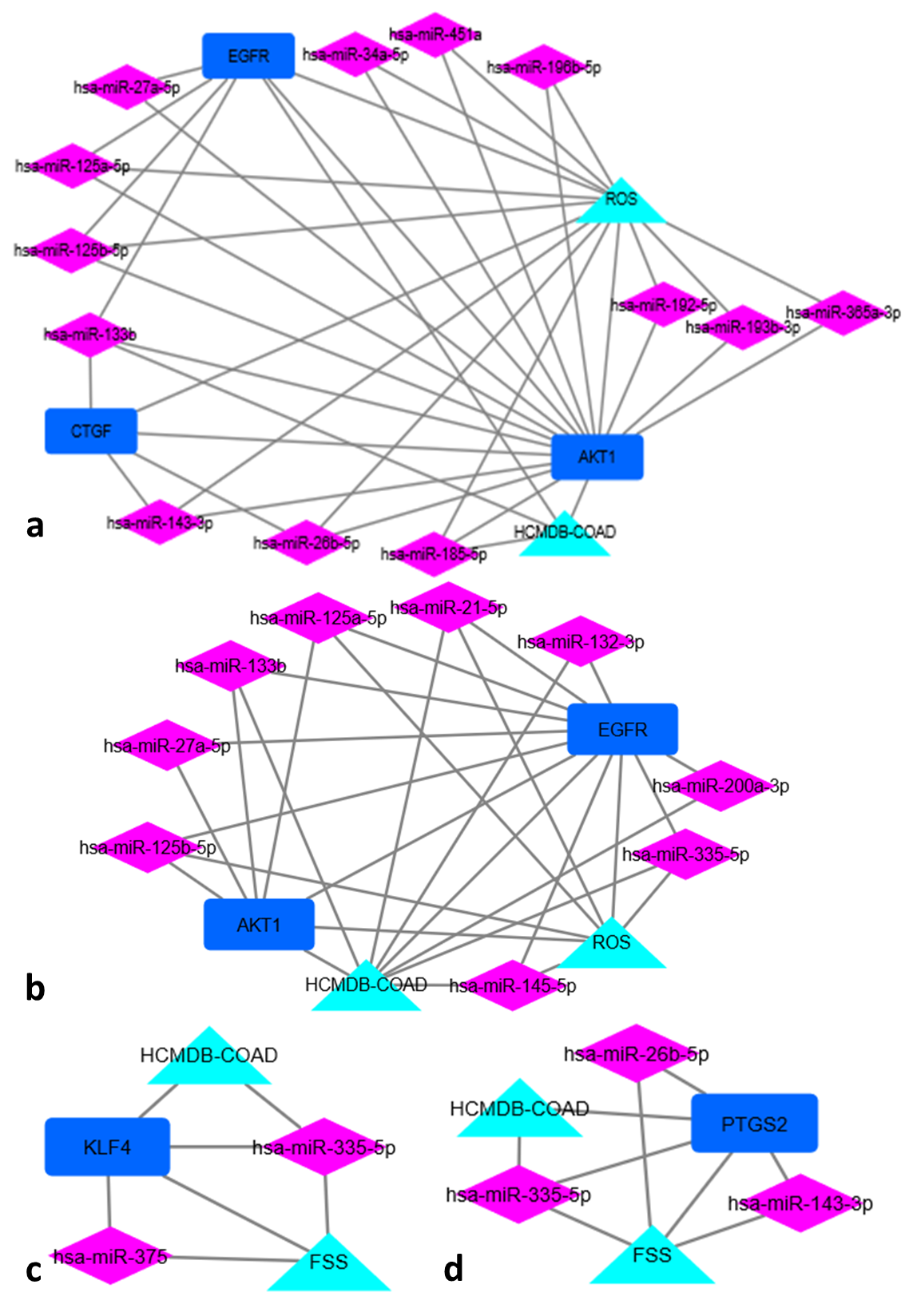
**Supplementary figure 7: Module analysis using FRM mRNA-mRNA-miRNA network as the input: Extraction of modules showing direct association of identified miRNAs with either ROS and HCMDB-COAD (Supp. fig. 7a and 7b) or FSS and HCMDB-COAD (Supp. fig. 7c and 7d) [FRM – FSS, ROS and HCMDB-COAD]**

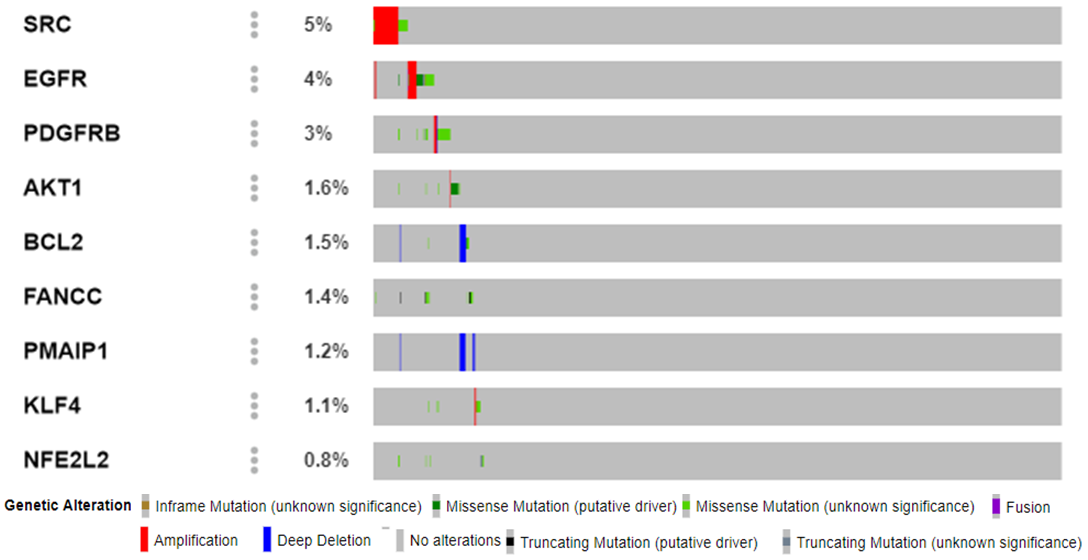

**Supplementary figure 8: Oncoprint analysis of nine significant mRNAs associated with FSS, ROS, and HCMDB-COAD obtained using MSKCC metastasis data.**
